# CD62L-selected umbilical cord blood universal CAR T cells

**DOI:** 10.1101/2024.01.18.576145

**Authors:** Christos Georgiadis, Lauren Nickolay, Farhatullah Syed, Hong Zhan, Soragia Athina Gkazi, Annie Etuk, Ulrike Abramowski-Mock, Roland Preece, Piotr Cuber, Stuart Adams, Giorgio Ottaviano, Waseem Qasim

**Affiliations:** UCL Great Ormond Street Institute of Child Health, WC1N 1DZ, London, UK; Great Ormond Street Hospital for Children NHS Trust, WC1N 3JH, London, UK

## Abstract

Umbilical cord blood (UCB) T cells exhibit distinct naïve ontogenetic profiles and may be an attractive source of starting cells for the production of chimeric antigen receptor (CAR) T cells. Pre-selection of UCB-T cells on the basis of CD62L expression was investigated as part of a machine-based manufacturing process, incorporating lentiviral transduction, CRISPR-Cas9 editing, T-cell expansion and depletion of residual TCRαβ T cells. This provided stringent mitigation against the risk of graft versus host disease (GVHD), and was combined with simultaneous knockout of CD52 to enable persistence of edited T cells in combination with preparative lymphodepletion using Alemtuzumab. Under compliant manufacturing conditions, two cell banks were generated with high levels of CAR19 expression and minimal carriage of TCRαβ T cells. Sufficient cells were cryopreserved in dose-banded aliquots at the end of each campaign to treat dozens of potential recipients. Molecular characterisation captured vector integration sites and CRISPR editing signatures and functional studies, including *in vivo* potency studies in humanised mice, confirmed anti-leukaemic activity comparable to peripheral blood-derived universal CAR19 T cells. Machine manufactured UCB derived T cells banks offer an alternative to autologous cell therapies and could help widen access to CAR T cells.

## Introduction

Chimeric antigen receptor (CAR) T cells can mediate leukaemic remission in patients that have otherwise failed conventional treatment ^1^. Current authorised products are manufactured from autologous peripheral blood lymphocytes derived from steady-state apheresis, and this requires complex logistics, and time for manufacturing and release of products ^2^. The quality and fitness of such products is highly dependent on individual patient variables ^3^. Alternative allogeneic sources are being investigated as a starting material, including T cells from HLA matched haematopoietic stem cell transplant donors ^4, 5^, or donor derived virus specific T cells ^6^ and non-matched genome-edited donor derived T cells ^7-10^. The latter have been derived from adult volunteer donors and edited to try and address HLA barriers. Steps have included targeted disruption of TCRαβ expression in combination with HLA class I knockout, alone or in combination with class II inhibition ^11-16^. We have previously tested strategies to confer resistance to lymphodepleting serotherapy by editing of CD52 and showed that cells persisted for around four weeks in the presence of Alemtuzumab, sufficient time to mediate leukaemic clearance ^8, 9, 17^.

Umbilical cord blood (UCB) T cells have distinct ontogenetic origins and may offer enhanced properties of expansion and activation, compared to adult peripheral blood lymphocytes ^18^. The specific transcriptional signature of UCB T cells has been affiliated with potent anti-leukaemic effects after allogeneic transplantation ^19^. Several groups have explored the efficacy of UCB-derived immune cells modified to express CARs in pre-clinical models, and clinical testing of cord derived natural killer (NK) cells has been underway in patients with B-cell malignancies, with encouraging early data ^20-23^. UCB T-cell naivety and high proliferative capacity warrants their further investigation for CAR T cell production ^24^. Challenges include the limited volume of UCB collections and their high nucleated red cell and mononuclear cell content which complicates T cell isolation and processing. CD62L is expressed on almost all UCB T cells and we reasoned that selecting for CD62L could allow T cells to be efficiently isolated from umbilical collections. CD62L generally identifies less-differentiated T cells (naïve and central memory T cells) from effector subsets ^25, 26^ and represents an important homing marker ^27^. Importantly, selection by targeting CD62L avoids binding of T-cell receptors or other key activation ligands, and this may be important for downstream activation and transduction steps. T cells are most efficiently transduced while undergoing mitosis and this is generally best achieved through a combination of anti-CD3 and anti-CD28 antibody stimulation. Previously we generated adult donor CAR19 T cells, with T-cell receptor alpha constant gene (*TRAC*) and *CD52* multiplexed knockouts using lentiviral gene modification of peripheral blood CAR T cells for a Phase I clinical trial in children with refractory/relapsed (R/R) B-ALL ^8^. These processes were now adopted for ‘compliance-ready’ machine manufacture of UCB T cells, starting with CD62L positive selection and ending with TCRαβ negative selection of TCRαβ depleted CAR19 T cells. Molecular, phenotypic and functional assessments were undertaken to determine suitability and feasibility of downstream therapeutic applications.

## Materials and Methods

### Manufacture of UCB-TT52CAR19

Fresh umbilical cord blood units were collected under ethical approval from volunteers identified independently by the Nolan transplant registry and tissue typed at St Barts Hospital National Health System (NHS) Trust. A unit was directly attached to the CliniMACS Prodigy device (Miltenyi Biotec). T cells were enriched by automatic red blood cell depletion and CD62L positive selection. The CD62L^+^ UCB T cells were then activated using MACS GMP TransAct (Miltenyi Biotec) on a modified T-cell transduction (TCT) program and cultured in TexMACS supplemented with 3% human serum (HS) (Life Science Production) and interleukin-2 (IL-2) (Miltenyi Biotec) for 24 hours. The cells were transduced on device with TT52CAR19 lentiviral vector, previously described ^8^. On day 4 post activation, cells were removed from the CliniMACS Prodigy and electroporated with capped, polyadenylated, and uridine modified SpCas9 mRNA using the Lonza 4D-Nucleofector. Post electroporation, cells were returned to the CliniMACS Prodigy and cultured at 5% CO_2_ 37°C until day 11 with timed TexMACS supplemented with 3% HS/IL-2. Programmed media changes with shaking enabled for optimal gas exchange. On day 11 cells underwent automatic TCRαβ depletion on the CliniMACS Prodigy using an anti-biotin bead kit. The depleted cells were rested overnight in the device and the final drug substance was harvested on day 12 and cryopreserved in 1 x 10^7^ and 2 x 10^7^ doses.

## Phenotype, Function and Molecular Characterisation

### Flow cytometry

Flow cytometry was undertaken using a BD FACSCanto II (Becton, Dickinson BD Biosciences) at Great Ormond Street Hospital (GOSH) NHS Trust, with additional characterisation using a BD LSRII (Becton, Dickinson BD Biosciences), and analysis using FlowJo v10 (TreeStar Inc.).

### Quantification of on- and off-target editing effects and translocations

PCR amplicons of target genomic DNA were Sanger sequenced and non-homologous end joining (NHEJ) events analysed using Inference of CRISPR Edits (ICE) protocols (https://ice.synthego.com/#/). On-target and off-target sites were informed by previous Digenome-seq studies ^7, 28^ **(Table S1)** and libraries subjected to paired-end Next Generation Sequencing (NGS) using MiniSeq (Illumina) as previously described ^29^. Droplet digital PCR (ddPCR) was used to quantify predicted translocations between chromosomes 14q (TRAC) and 1p (CD52) loci and analysed using Quantasoft (BioRad) **(Table S1)**. (BioProject accession number: PRJNA1061060)

#### *In vitro* cytotoxicity assay

*In vitro* cytotoxic function of UCB-TT52CAR19 GMP1 (UCB1) and UCB-TT52CAR19 GMP2 (UCB2) or peripheral blood leukapheresis (PBL) derived batches alongside controls was quantified by co-culture with chromium (^51^Cr) loaded CD19^+^ Daudi and CD19^+^ or CD19^-^ SupT1 cells for 4 hours at 37°C at increasing effector-to-target (E:T) ratios and ^51^Cr release was quantified using a Wallac MicroBeta TriLux microplate scintillation counter.

#### *In vivo* studies

*In vivo* function was assessed in non-obese diabetic (NOD)/severe combined immunodeficiency (SCID)/γc (NSG) mice inoculated intravenously by tail vein injection with 0.5 x 10^6^ CD19^+^ Daudi tumour cells expressing enhanced green fluorescence protein (eGFP) and luciferase, followed by 5 x 10^6^ UCB1, UCB2 or PBL derived effectors and non-edited or non-transduced controls after 4 days. Serial bioluminescence imaging using an IVIS Lumina III In Vivo Imaging System (PerkinElmer, Living Image® version 4.5.5) was used to track leukaemia inhibition for up to 4 weeks.

### Statistical analysis

Statistical analyses were performed with Prism (GraphPad) using unpaired t test or ANOVA where indicated. Values from 3 or more samples are presented as mean with SEM. P < 0.05 was considered significant.

## Results

### Machine mediated CD62L^+^ T-cell enrichment and engineering of UCB donations

Banks of UCB-TT52CAR19 (UCB1 and UCB2) were manufactured from fresh cord blood donations using a CliniMACS Prodigy device **(Figure 1)**. One key hurdle in handling and manipulating UCB donations is the high number of nucleated red cells. We investigated CD62L^+^ T-cell enrichment by CD62L selection using magnetic bead positive selection. A yield of >4 x 10^7^ CD62L^+^ cells was set as a starting material minimum before activation with anti-CD3/CD28 Transact reagent and lentiviral transduction. We found that this was readily achieved, with the umbilical cord blood units yielding: 1.33 x 10^8^ (UCB1) and 0.96 x 10^8^ (UCB2) from donations of 4.4 x 10^8^ and 4.2 x 10^8^ MNCs, respectively (**Figure 2A, B)**. Using MOIs of around 5, transduction efficiencies of 67% (UCB1) and 74% (UCB2) were achieved (**Figure 2C; Figure S1**).

**Figure 1.**
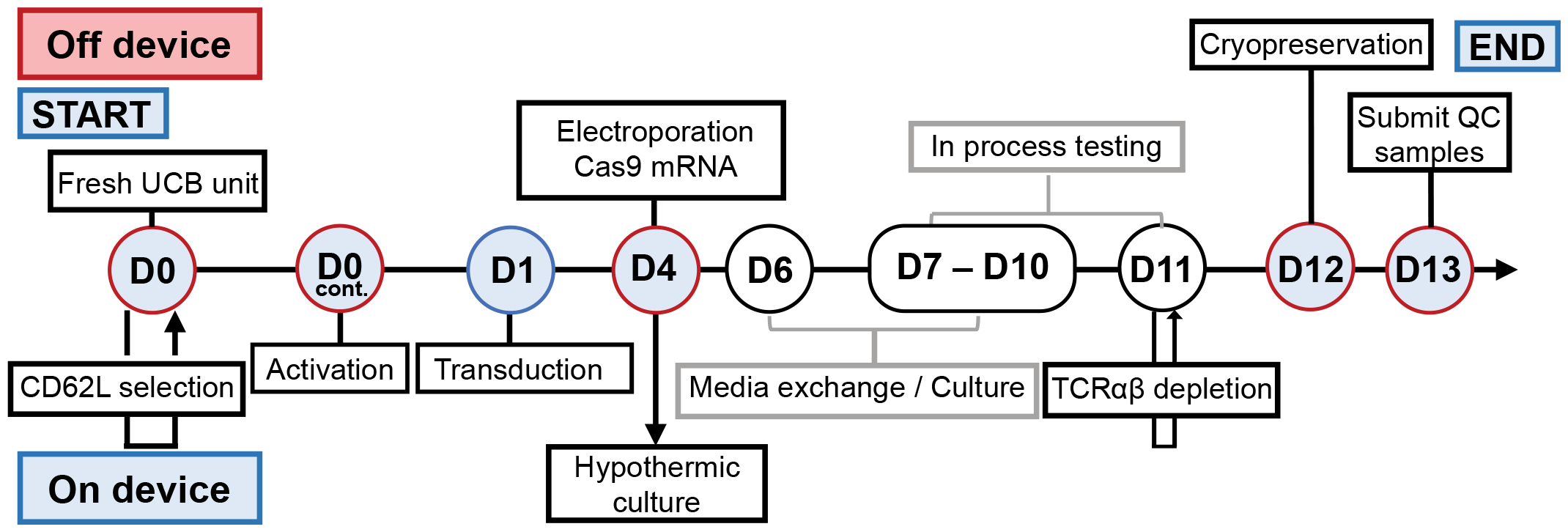
Schematic timeline for GMP manufacturing of TT52CAR19 cell banks from Umbilical Cord Blood (UCB) starting material. Fresh UCB units were enriched for lymphocytes by CD62L^+^ magnetic bead positive selection and activated on day 0 (D0). T cells were transduced 24hrs later (D1) with TT52CAR19 vector and electroporated on day 4 (D4) with SpCas9 mRNA. On day 11 (D11), cells underwent automated TCRαβ T-cell depletion. After this step the final cell product was harvested and cryopreserved in therapeutic doses. Steps itemised below the timeline were performed on a Miltenyi Prodigy and are shown as ‘On-device’, with the remaining items (above the timeline) undertaken off-device.

**Figure 2.**
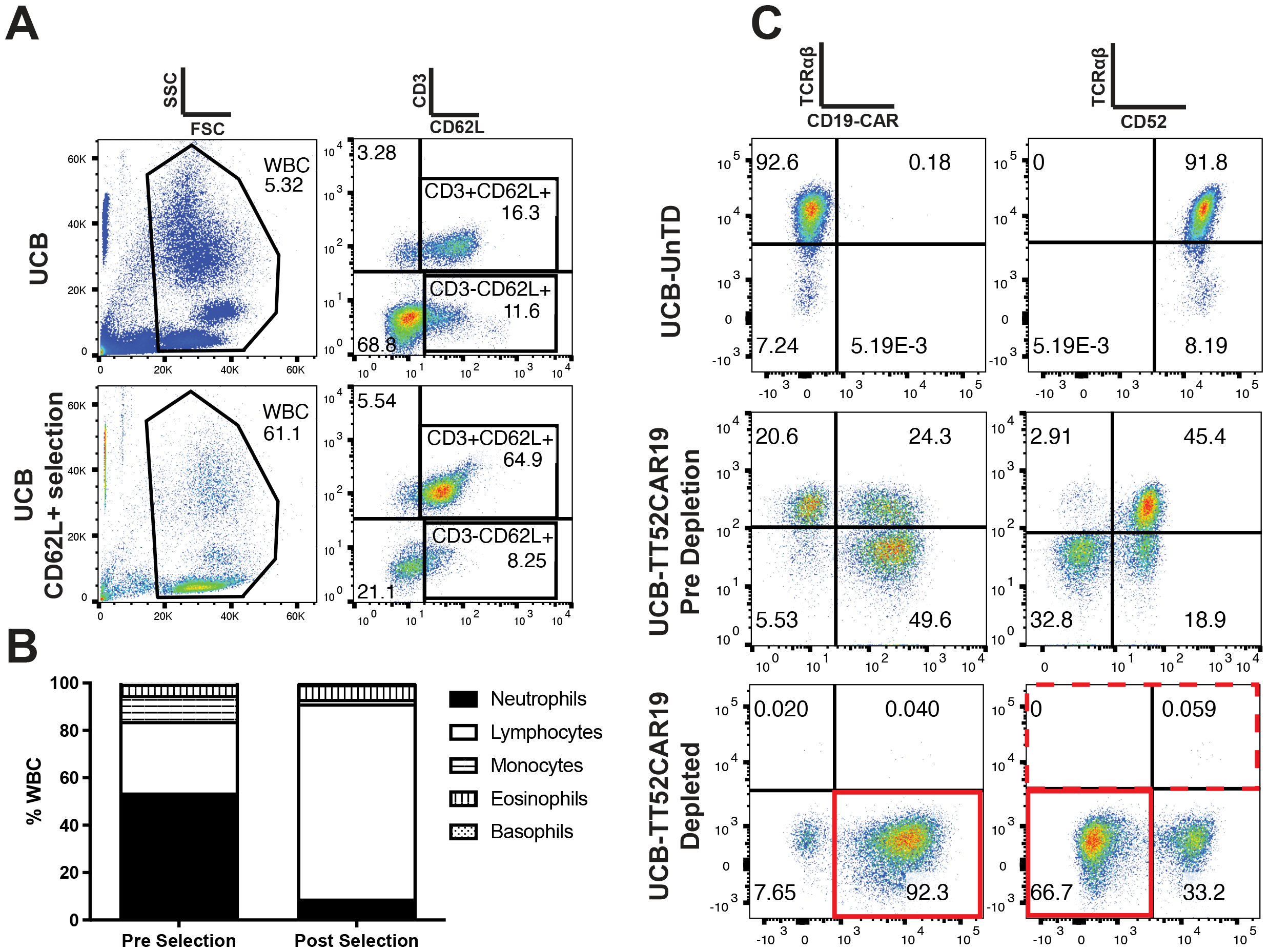
Whole UCB CD62L selection using the CliniMACS Prodigy and TT52CAR19 batch manufacture. **A**. UCB cells pre and post CD62L^+^ selection using the CliniMACS Prodigy demonstrates enrichment of CD3^+^ T cells. **B**. Cord blood cells from UCB donors (n=4) were analysed by Sysmex pre and post CD62L selection using the CliniMACS Prodigy. **C**. Flow cytometric T-cell transduction and knockout analysis of UCB-TT52CAR19 GMP1 (UCB1) cell bank. UCB-TT52CAR19 cells were stained before and after up to two rounds of magnetic TCRαβ depletion alongside untransduced (UnTD) cells or non-edited UCB-TT52CAR19 TCR^+^ cells. Transduction efficiency was measured by quantifying transgene expression using F(ab’)2, and CRISPR-Cas9 mediated protein knockout was determined through staining for TCRαβ and CD52.

Next, electroporation was performed ‘off-device’ using a Lonza LV for transient delivery of SpCas9 mRNA. Flow cytometry on day 7 found residual TCRαβ^+^ (UCB1 34.6%; UCB2 35.4%) and CD52^+^ (UCB1: 35.4%; UCB2 30.2%) (**Figure 2C; Figure S1**). ‘On-device’ prodigy expansion for 3 days was followed by automated bead-mediated depletion of residual TCRαβ T cells with UCB1 exhibiting 0.2% and UCB2 0.3% TCRαβ cells at the end of processing. Simultaneously, the CAR^+^ fraction increased to >80% by the end of production as a result of coupling effects of editing and transduction in the TT52CAR19 configuration. Yields of 9.1 x 10^8^ and 5.5 x 10^8^ cells in total were achieved for UCB1 and UCB2, respectively. The product was cryopreserved in 1 ml aliquots of 1 x 10^7^ (x 20 vials for each UCB1 and UCB2 batch) and 2 x 10^7^ (x 10 vials for each UCB1 and UCB2 batch) total cells in individual vials and stored at <-130 °C **(Table 1)**. Western blot and ELISA did not detect presence of Cas9 at the end of production **(Figure S2)**.

**Table 1.** UCB-TT52CAR19 GMP1 and GMP2 end of manufacture batch specifications.

|  | UCB-TT52CAR19-GMP1 (UCB1) | UCB-TT52CAR19-GMP2 (UCB2) |
| --- | --- | --- |
| **Viability (7AAD)** | 96.2% | 99.8% |
| **CAR19 Transduction Efficiency** | 88.7% | 82.8% |
| **Purification** | 0.2% CD45+TCRαβ+ cells | 0.3% CD45+TCRαβ+ cells |
| **Sterility (BACTEC)** | No Growth | No Growth |
| **Sterility (Gram stain)** | No Organisms Seen | No Organisms Seen |
| **Mycoplasma** | None Detected | None Detected |
| **Endotoxin** | <1.0 EU/ml | <1.0 EU/ml |
| **Yield** | 906 x 10<sup>6</sup> | 546 x 10<sup>6</sup> |

### Phenotype & Molecular Characterisation of UCB-TT52CAR19

At the end of production >80% of cells were CD45RA^+^CD62L^+^ (**Figure S3)** with 85% - 92% CAR expression. Transduction was corroborated by proviral copy number by qPCR (UCB1 VCN: 2.7; UCB2 VCN: 4.0 copies/cell). On-target genome editing signatures of NHEJ were verified by quantification of insertions/deletions (indels) at TRAC (UCB1: 76%; UCB2: 59%) and CD52 (UCB1: 58%; UCB2: 51%) loci by ICE analysis of Sanger sequence traces (**Figure 3A; Figure S4A**). Targeted ddPCR quantification for NHEJ at these sites provided corroboration (TRAC: 65%, 47%; CD52: 73%, 38%) (**Figure 3B; Figure S4B**). Lentiviral integration was mapped by LM-PCR with the top ten most frequent sites presented in **Figure 4; Figure S5**.

**Figure 3.**
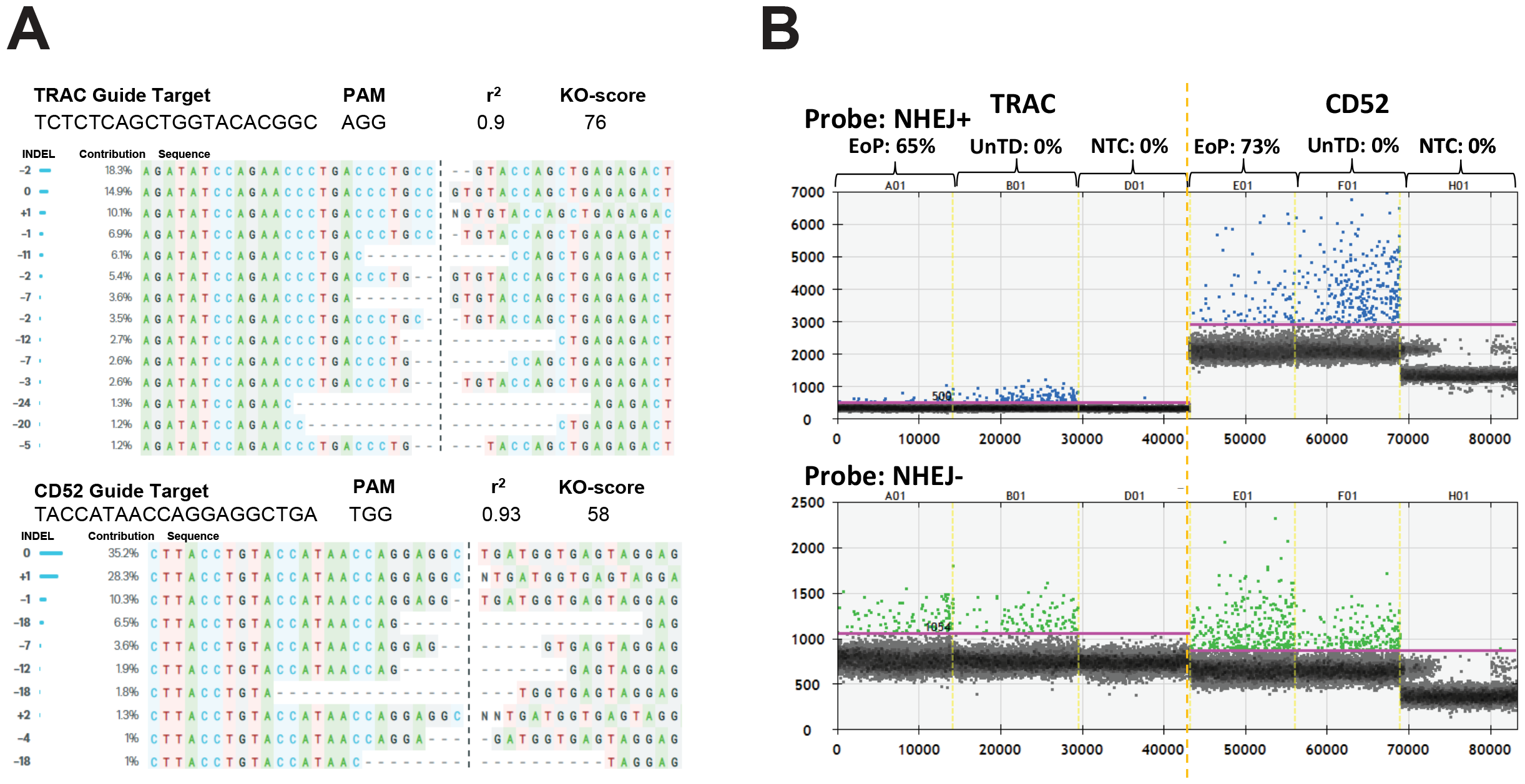
Molecular characterisation of on-target CRISPR-Cas9 mediated cleavage. **(A)** ICE analysis of Sanger sequence traces identified indels as signatures of NHEJ at the TRAC locus and at the CD52 target site. **(B)** Quantification of indels by ddPCR at both the TRAC and CD52 sites using separate probes, one specific to the predicted NHEJ region (NHEJ^+^) and a second outside the NHEJ region (NHEJ^-^).

**Figure 4.**
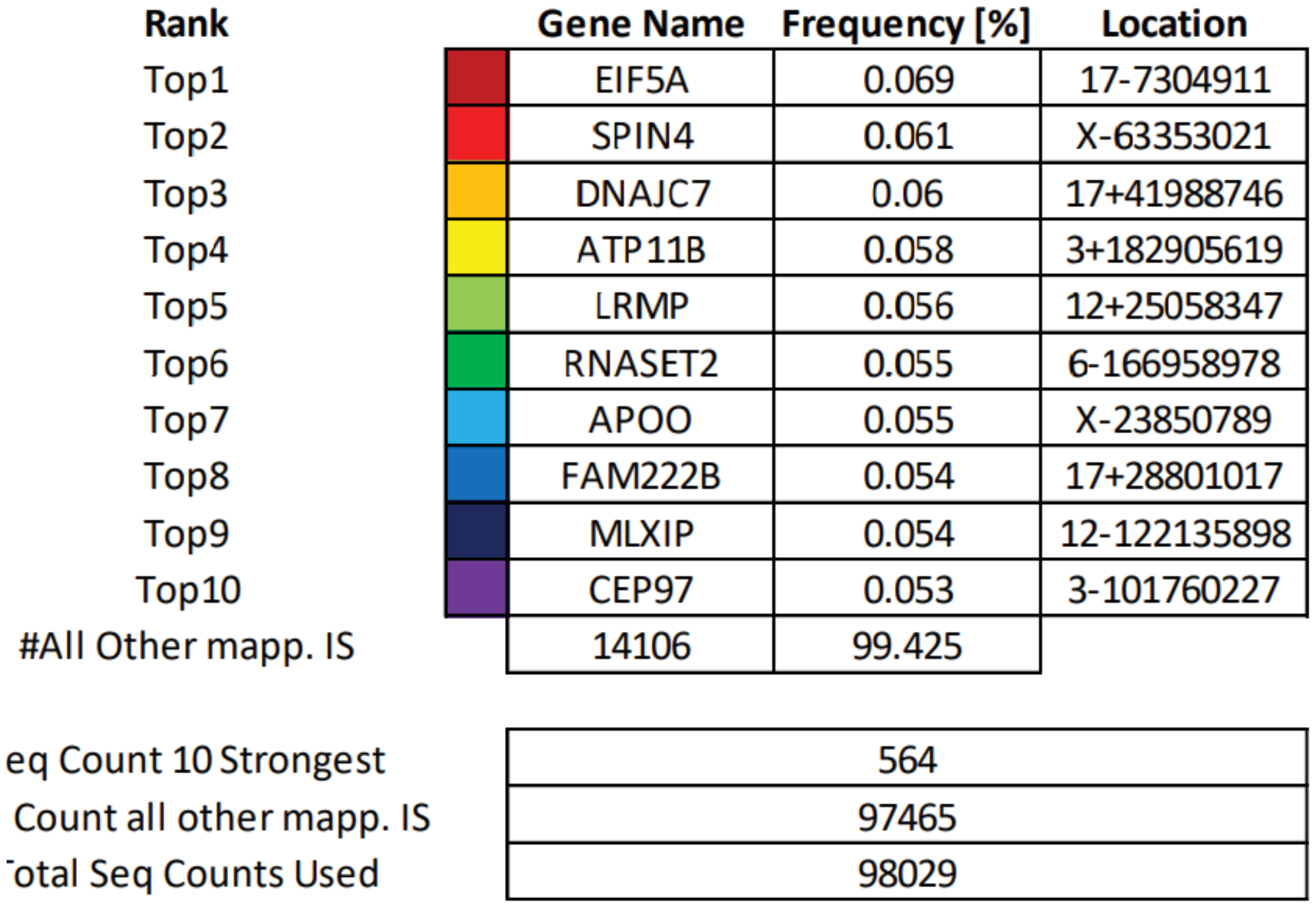
Integration site analysis. Ligation-mediated PCR (LMPCR) detection and quantification of vector integration sites (IS) where the top 10 most frequent sites comprised <0.1% of integrants.

### End of production screening for aberrant DNA breaks

Karyotype analysis reported normal G-band analysis with no detection of chromosomal aberrations. Further investigations by FISH analysis found no evidence (within the limits of the probe set) of TRAD rearrangements in 197/200 cells (98.5%). Predictable translocation events were investigated (**Figure 5A; Figure S6A**) for four predicted recombinations (C1-C4) involving 14p (TRAC locus) and 1q (CD52 locus), and low frequency events (<1.0%) were quantified by ddPCR for each reaction, with 0.64% (UCB1) and 0.73% (UCB2) in total quantified over the four reactions.

**Figure 5.**
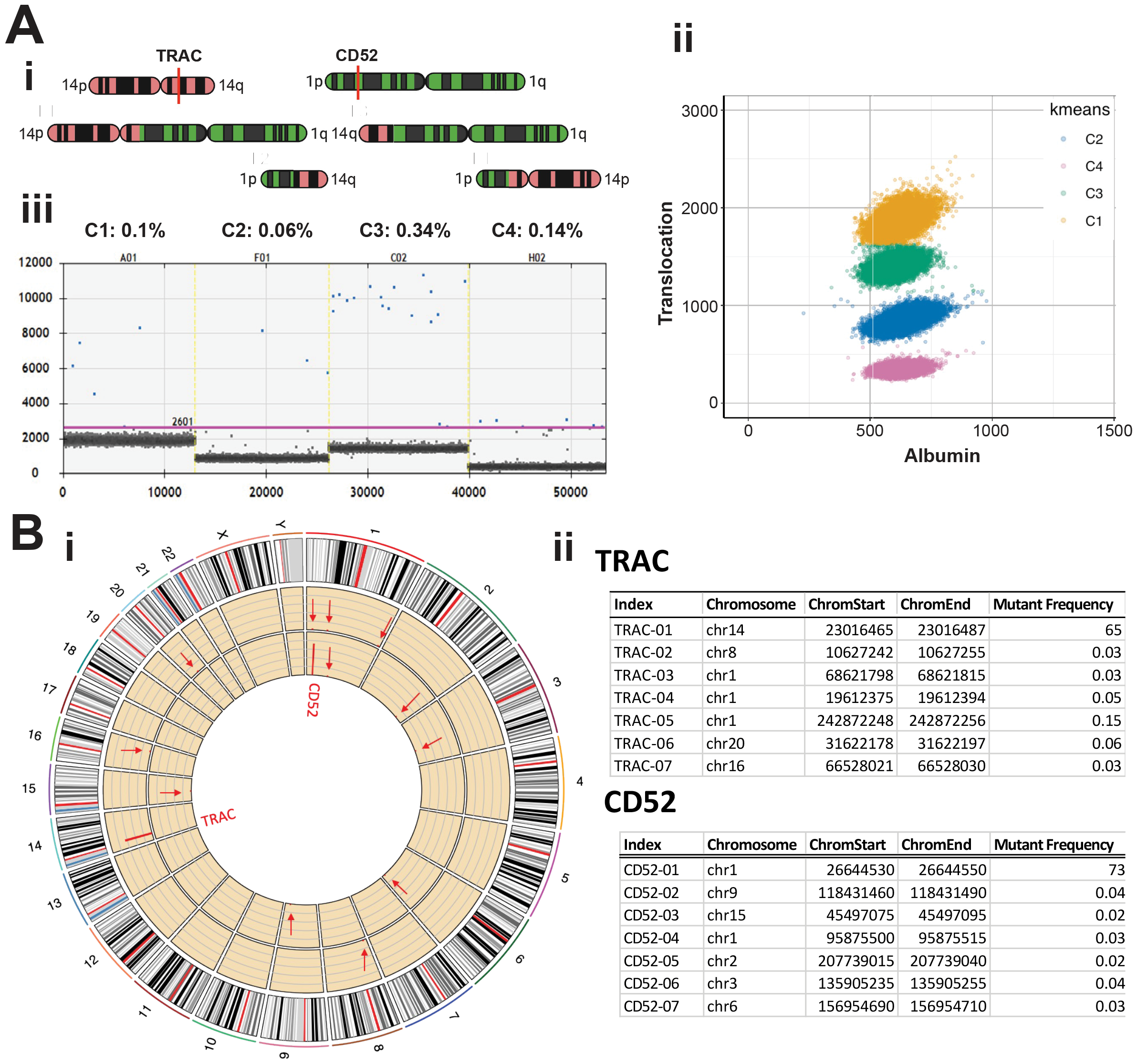
Detection for translocation and off-target CRISPR-Cas9 mediated cleavage events. **(A)** Droplet Digital PCR (ddPCR) was used for the detection and quantification of possible translocations after ‘on-target’ DNA scission. Four predicted recombination events (C1-C4) are presented in the schematic with TRAC (Chr14q) (green), CD52 (Chr1p) (red) and SpCas9 cleavage site (red line). Right panel colours (yellow, green, magenta, blue) discriminate four possible translocations. Left panel shows low frequency translocation events (blue dots) C1-C4 arising between the edited TRAC and CD52 loci. Cumulative events for all four possible events were <1%. **(B)** Circos plot with verification of quantification using targeted NGS across six highest scoring predicted off-target sites. TRAC-01 (solid red line marking locus in outer yellow circle), CD52-01 (solid red line marking inner yellow circle) and at predicted off-target sites TRAC-02-TRAC-07 (red arrows marking outer yellow circle) and CD52-02-CD52-07 (red arrows marking inner yellow circle). Table shows that negligible off-target events were detected for both the TRAC and CD52 guides.

Previously, Digenome-Seq analysis had informed design of screening for ‘off-target’ guide-dependent editing for the TT52CAR19 vector system. An abbreviated form of the screen was now applied at the six highest scoring genomic sites (**Figure 5B; Figure S6B**). Targeted NGS found modification frequencies at these sites of <0.5% in both batches, whereas on-target NHEJ events were quantified as 65% and 51% at the TRAC locus and 73% and 17% at the CD52 locus for UCB1 and UCB2, respectively.

### Functional studies of UCB-TT52CAR19 cells

The potency of each UCB-TT52CAR19 batch was assessed against ^51^Cr labelled CD19^+^ Daudi cells and CD19^+^ or CD19^-^ SupT1 cells (**Figure 6A; Figure S7A-C**). Specific lysis was demonstrated across a range of E:T ratios compared to untransduced (UnTD) cells (**P<0.01) with results comparable to TT52CAR19 products generated from adult PBL (**Figure 6B**). In addition, cytokine production was measured following a 1:1 co-culture with CD19^+^ Daudi cells and CD19^+^ or CD19^-^ SupT1 cells (**Figure 6C; Figure S7D-F**) and secretion of cytokines IFN-γ, TNF-α, IL-4 and IL-2, compared to PBL derived TT52CAR19 cells (**Figure 6D**).

**Figure 6.**
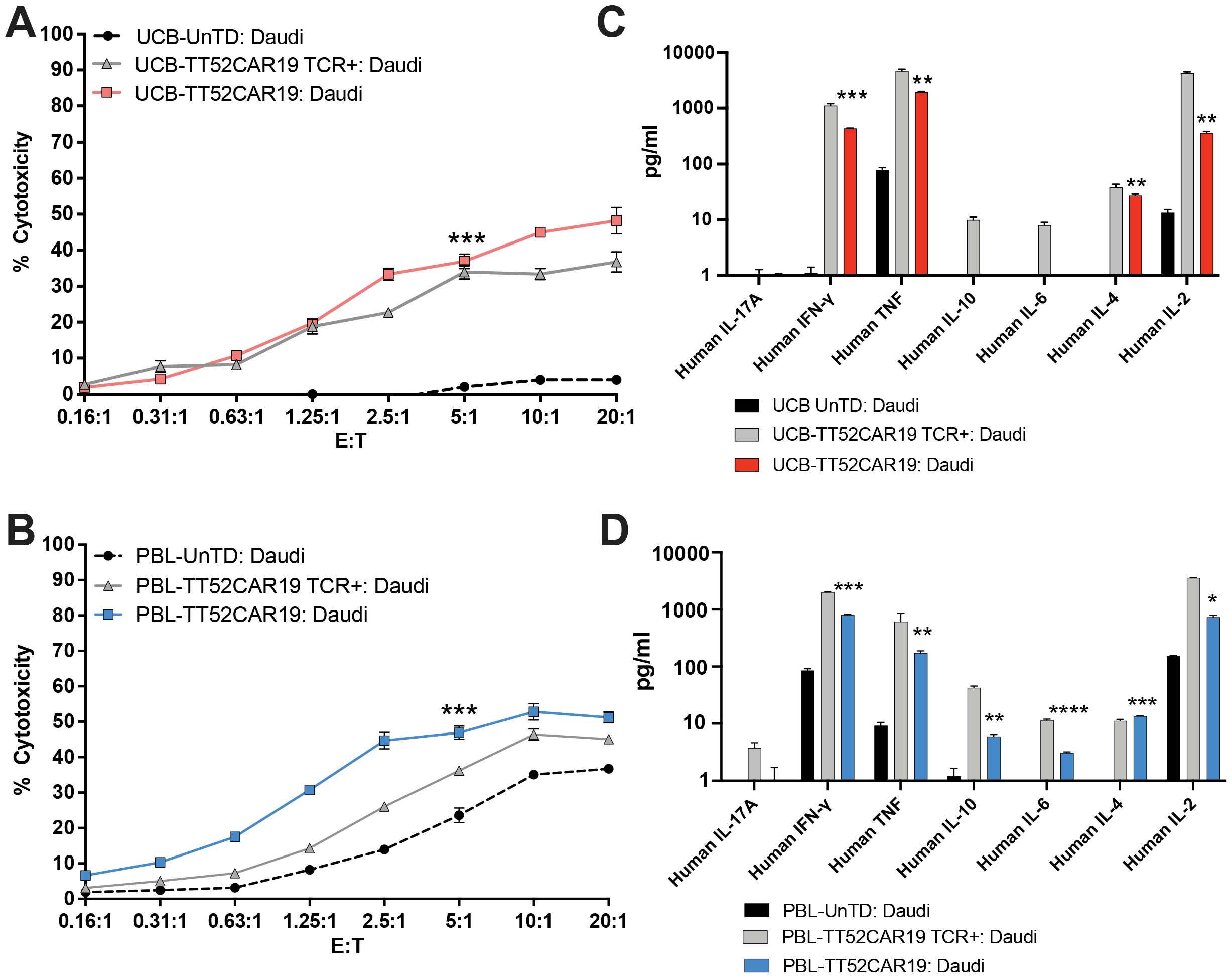
*In vitro* killing potential and cytokine production of UCB-TT52CAR19 cells against CD19^+^ targets. *In vitro* cytotoxicity of UCB-TT52CAR19 GMP1 (UCB1) cell bank compared to respective UCB-TT52CAR19 transduced but not edited (UCB-TT52CAR19 TCR^+^) or non-transduced (UnTD) controls **(A)** or PBL-TT52CAR19 effectors and respective controls **(B)** when measured by ^51^Cr chromium release of labelled CD19^+^ Daudi cells following four hours of co-culture at incremental effector (E) – target (T) ratio. Effector responses were considered successful if ≥50% lysis was detected compared to UnTD controls at E:T cell ratios between 1.25:1 and 5:1. Error bars SEM, n=3 replicate/wells. *P < 0.05, **P < 0.01, ***P < 0.001 by t-test. In vitro target specific cytokine secretion of UCB-TT52CAR19 GMP1 cell bank (UCB1), and respective UCB-TT52CAR19 TCR^+^ or non-transduced (UnTD) controls after co-culture with CD19^+^ Daudi cells overnight **(C)**. Cytokine release also quantified for PBL-TT52CAR19 effectors and respective PBL-TT52CAR19 TCR^+^ and UnTD control co-cultures with CD19^+^ Daudi cells **(D)** The presence of cytokines in the co-culture supernatant was measured by cytokine bead array and levels >50 pg/ml were considered positive responses. Significance was calculated between UCB-TT52CAR19 GMP1 (UCB1) or PBL-TT52CAR19 banks and respective UnTD controls. *P < 0.05 by t-test. Error bars represent SEM, n=3 replicates.

A chimeric human:murine xenograft model of B-cell malignancy was used to assess *in vivo* function of UCB-TT52CAR19 cells (**Figure 7A**). NSG mice were engrafted with 5 x 10^5^ CD19^+^eGFP^+^Luciferase^+^ Daudi cells and 5 x 10^6^ UCB-TT52CAR19 effectors were infused 4 days later. Serial bioluminescence imaging over a 4-week period showed rapid disease progression in mice receiving UnTD cells, whereas mice treated with UCB-TT52CAR19 cells exhibited significantly reduced tumour signal and inhibition of disease throughout the monitoring period (UCB-UnTD GMP1 vs UCB-TT52CAR19 GMP1 (UCB1): P <0.0001; UCB-UnTD GMP2 vs UCB-TT52CAR19 GMP2 (UCB2): P <0.0001) (**Figure 7B, C; Figure S8**). Anti-leukaemic activity was similar to animals treated with PBL TT52CAR19 effector batches (UCB1 vs PBL P= 0.975; UCB2 vs PBL P = 0.986).

**Figure 7.**
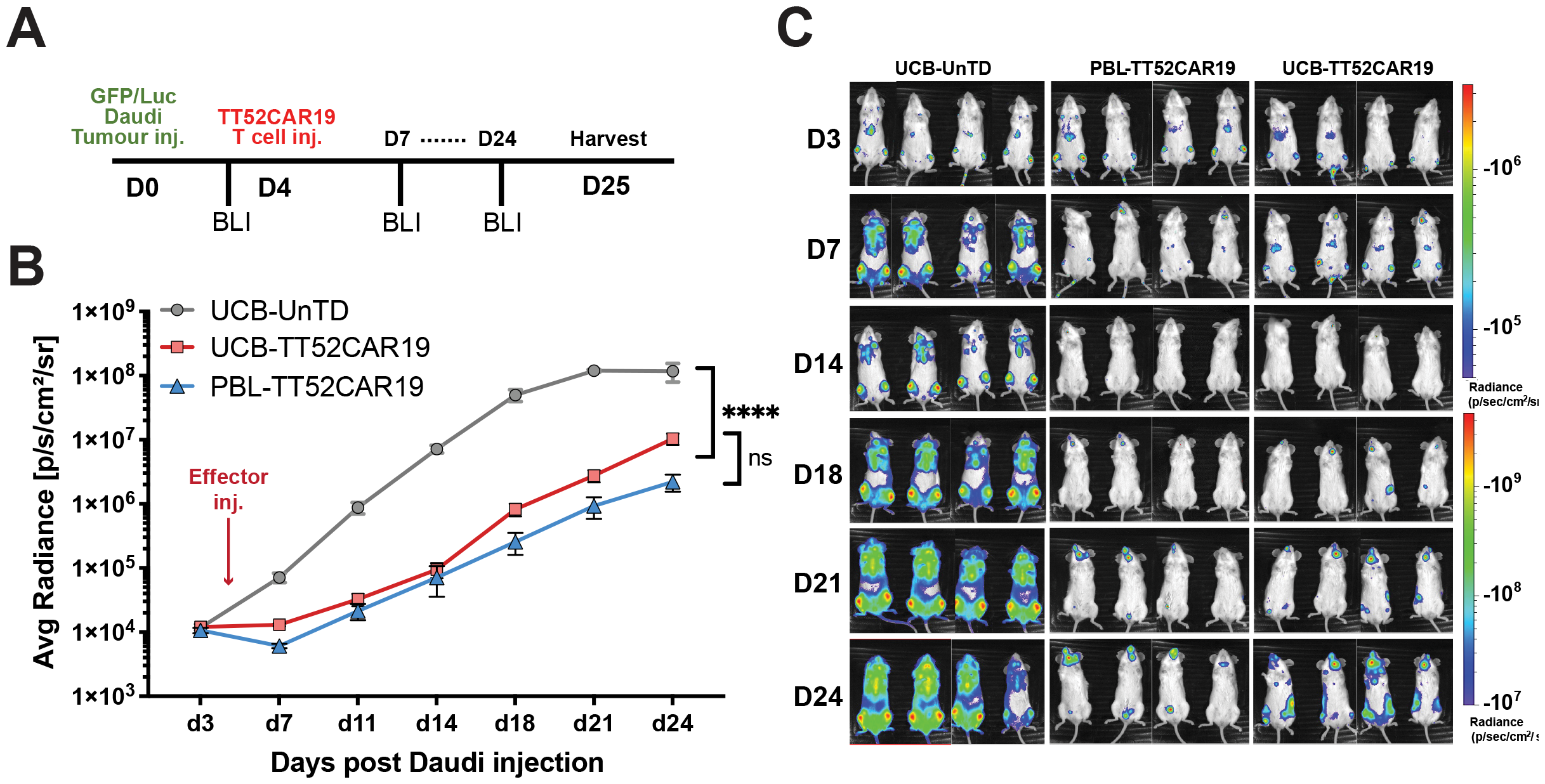
*In vivo* tumour clearance in human:murine xenograft model of CD19^+^ disease. Timeline of *in vivo* human:murine xenograft modelling indicating target and effector intravenous injection days and bioluminescent imaging (BLI) timepoints **(A)**. Serial measurement of bioluminescence of Daudi CD19^+^ B-cell disease in immunodeficient mice NOD/SCID/γc (NSG) mice (n=12) infused with GFP/luciferase expressing Daudi CD19^+^ B cells were treated on day 4 with either umbilical cord blood (UCB) UCB-TT52CAR19 GMP1 (UCB1) (n=5) or peripheral blood lymphocyte (PBL) PBL-TT52CAR19 (n=4) and were monitored over a 4 week period **(B, C)** Non-transduced (UnTD) (n=4) T cells were used as controls. Error bars represent SEM. Significance compared by area under the curve using one-way ANOVA (F-value 35.86) with Tukey multiple comparison post-hoc; not significant (ns) P ≥ 0.05; ****P < 0.0001.

## Discussion

Despite breakthroughs using autologous CAR T cells, there are notable hurdles to wider access to CAR therapies. Alternative “off-the-shelf” CAR T-cell banks suitable for multiple patients could ultimately reduce costs and avoid delays. Donor peripheral blood T cells have been successfully used to generate “universal” allogeneic CAR T cells. Umbilical cord donations may offer alternative sources of immune cells with advantageous immunological properties. UCB derived NK cells have been of interest for their capacity to be used without HLA matching ^22^. Trials have been underway for UCB NK cells transduced to express CAR19 and IL-15, to support antigen-driven expansion ^21^. In a phase I clinical trial (NCT03056339) the first patients exhibited anti-leukaemic activity without severe toxicities ^20^. Could cord blood T cells offer favourable expansion, persistence or longevity compared to adult PBLs? The importance of starting T-cell subsets for CAR-T manufacturing has been reported in pre-clinical and clinical studies ^30, 31^. When peripheral blood selected naïve/stem cell memory (N/SCM) CD62L-positive CAR T cells were compared to unselected T cells, they appeared to provide better expansion and persistence and anti-leukaemic responses in chimeric mice ^32, 33^. One important caveat has been the interpretation of flow-based phenotyping using conventional memory/naïve panels which may not be straightforward after activation and transduction with CARs with different activation domains. Previous studies have included UCB units transduced with a retroviral vector incorporating CAR19 and an IL-12 transgene, which exhibited central memory phenotypes by the end of manufacture and expression of cytotoxic effector proteins Granzyme B and IFNγ ^23^. Potent anti-leukaemic clearance in a preclinical mouse model of B-ALL was reported. In other work, cord blood units expressing anti-CD123 CAR T cells retained a less differentiated phenotype after activation and transduction, with anti-leukaemic activity *in vitro* and *in vivo* models ^34^. We reasoned that CD62L could be an ideal selection target of cord T cells for manufacturing CAR T cells. In addition, a key hurdle in manufacturing from cord blood involves distinguishing T cells from other mononuclear cells. Cord blood is rich in nucleated red cells, causing difficulties in enumeration and analysis, and creating challenges for setting optimal culture and transduction conditions. Pre-selection using anti-CD62L magnetic bead enrichment allowed efficient automated enrichment of cord T cells ahead of activation. The process could be readily incorporated into a CliniMACS Prodigy matrix, and with compliant reagents already available, included in established transduction and editing processes.

Overall, we found that CD62L^+^ cord CAR19 T cells exhibit expansion and anti-leukaemic activity *in vitro* and *in vivo*, comparable to control peripheral blood genome edited CAR19 T-cell products that have already been investigated in clinic ^8^. Cord collection offers a further advantage in allowing ready access to a broad range of HLA haplotypes with tolerance for mismatches. Ultimately, the risk of host mediated rejection could be mitigated using batches of cells generated from donors homozygous for common HLA haplotypes. This could help avoid the need for intensive lymphodepletion currently applied to address barriers caused by HLA mismatches. It has been estimated that a large proportion of European populations could be partly, or fully HLA matched,to ‘off the shelf’ cord blood banks derived from around 150 different donors ^35^.

In summary, UCB T cells can be isolated and enriched by CD62L selection and are amenable to machine-based gene modification, both using lentiviral vectors and by CRISPR-Cas9 editing. Potential therapeutic applications include ‘off the shelf’ CAR T therapies for ready access to products where autologous options are not available.

## Contributions

W.Q. is the principal investigator. C.G., G.O., L.N., F.S. and W.Q. wrote the first draft of the manuscript. C.G., A.E., R.P., and W.Q. developed the vector configuration and genome editing strategy and performed *in vitro* and *in vivo* phenotypic and functional assays. F.S., H.Z., and P.C. manufactured the UCB-TT52CAR19 batches and performed the longitudinal stability tests. S.A.G., and S.A. performed the molecular characterisation of the final product. C.G., G.O., S.A.G., and W.Q. analysed the data and created the figures of this manuscript. All authors reviewed and approved the manuscript.

## Acknowledgements

We would like to thank Ailsa Greppi and Kyle O’Sullivan for their invaluable technical support with *in vivo* studies. Dr. Ayad Eddaudi at the UCL Joint Great Ormond Street Institute of Child Health and Institute of Ophthalmology Flow Cytometry Core Facility, supported by the Great Ormond Street Children’s Charity (GOSHCC), grant reference U09822 (October 2007), UCL Capital Equipment Funding, School of Life and Medical Sciences (September 2012), and UK Research and Innovation, grant reference MR/ L012758/1 (March 2014). The Anthony Nolan Trust (803716/ SC038827) for their support and provision of healthy blood donations.

