## Supplementary Data for "CD62L-selected umbilical cord blood universal CAR T cells"

Supplementary Materials and Methods (page 2)
Supplementary figures (pages 3-10)
Supplementary Table (page 11)

#### Supplementary Materials and Methods

##### Determination of vector copy number (VCN)

CAR19 transgene in transduced cells was assessed by VCN in genomic DNA at end of manufacture of UCB-TT52CAR19 at GOSH NHS Trust. Vector copies were determined by qPCR or droplet digital PCR (ddPCR) targeting HIV-psi or human albumin sequences **(Table S1)**.

##### Karyotype and Fluorescence in Situ Hybridization (FISH)

UCB-TT52CAR19 cells were cultured with Colcemid overnight in order to arrest cells in metaphase. Karyotype using Giemsa-Banding (G-Banding) and fluorescence in situ hybridization (FISH) analysis was performed at GOSH NHS trust; the latter used a dual colour, break-apart FISH probe (Cytocell TCRAD LPH 047-S; 14q11, red/green fusion) to interrogate interphase nuclei.

##### Quantification of on- and off-target editing effects and translocations (continued)

On-target and off-target sites informed by previous Digenome-seq studies were captured using Phusion polymerase (New England BioLabs), and the products amplified using TruSeq HT Dual Index primers **(Table S1)**. Libraries were then subjected to paired-end Next Generation Sequencing (NGS) using MiniSeq (Illumina) as previously described. Demultiplexed fastq files were uploaded to Galaxy for quality checks, trimming and alignment. Non-homologous end joining (NHEJ) signatures were analysed using Pindel and figures were created in R.

##### Vector integration site analysis

Lentiviral integration sites (IS) were mapped by Genwerks (Protagene, Germany) using Shearing Extension Primer Tag Selection Ligation-Mediated PCR (S-EPTS/LM PCR) as previously described [1] and the top 10 most frequent loci tabulated.

##### Cytokine Bead Array

Cytokine release was quantified after overnight co-culture of UCB-TT52CAR19 GMP1 (UCB1) and GMP2 (UCB2) or PBL derived batches alongside TT52CAR19 TCR+ or UnTD control cells with CD19+Daudi and $CD19^{+}$ or $CD19^{-}$ SupT1 cells at a 1:1 ratio using a TH1/TH2/TH17 human bead array kit (Becton Dickinson Biosciences).

#### Supplementary Figure S1.

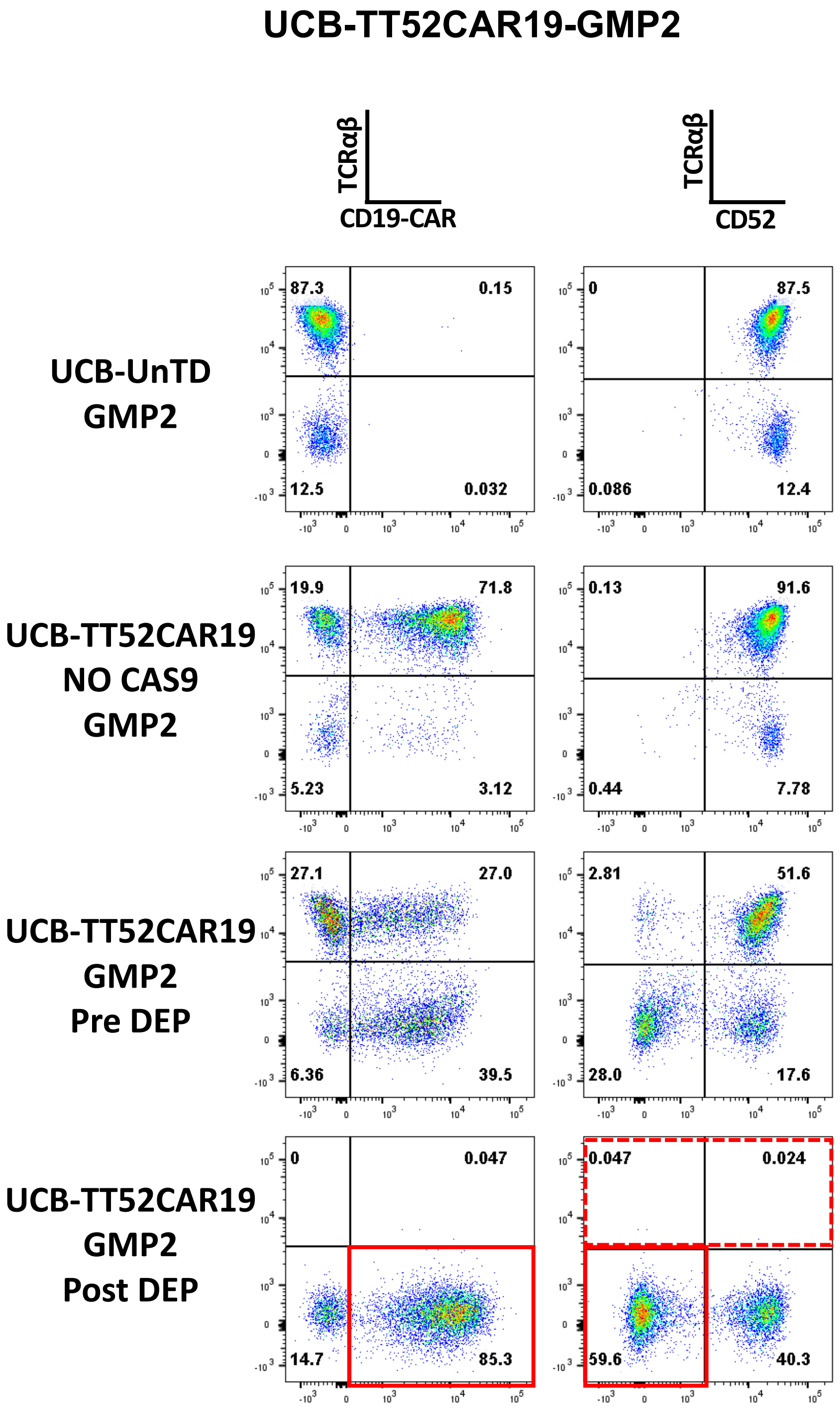


**Supplementary Figure S1. CAR19 and TCRαβ phenotyping of UCB-TT52CAR19 GMP2 bank throughout manufacture.** Flow cytometric T cell transduction and knockout analysis of UCB-TT52CAR19 GMP2 (UCB2) cell bank. UCB-TT52CAR19 cells were stained pre and post magnetic TCRαβ depletion alongside untransduced (UnTD) cells or non-edited UCB-TT52CAR19 TCR+ cells. Transduction efficiency was measured by quantifying transgene expression using F(ab')2, and CRISPR/Cas9 mediated protein knockout was determined through staining for TCRαβ and CD52.

#### Supplementary Figure S2.

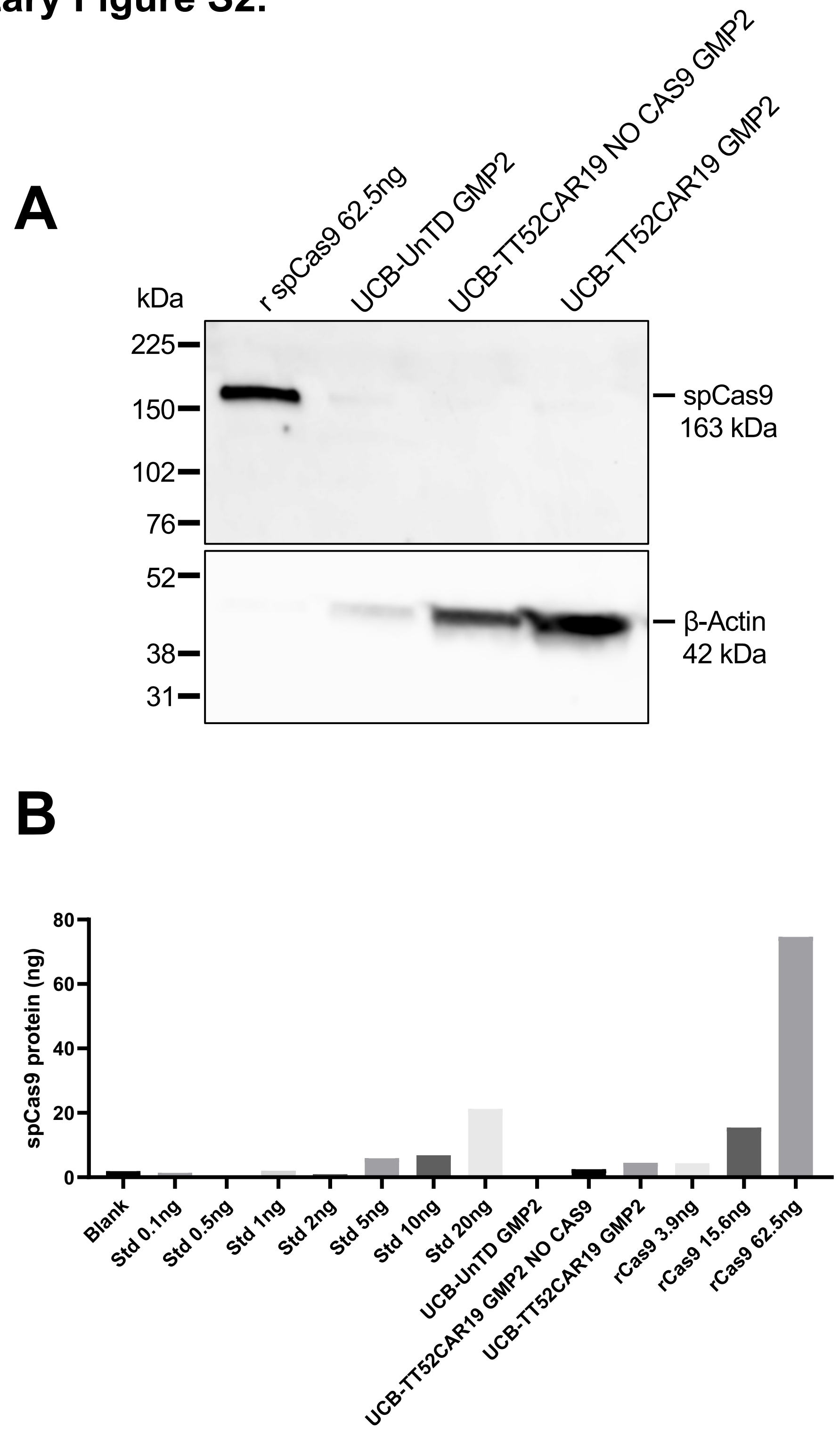


**Supplementary Figure S2. End of production verification of absence of residual SpCas9 protein. (A)** Whole cell lysate (20µg) from end of production UCB-TT52CAR19 GMP2 (UCB2) cells (lane 4), untransduced (UnTD) (lane 2) or non-edited UCB-TT52CAR19 NO CAS9 GMP2 (lane 3) was subjected to western blot SDS-PAGE and stained with a rabbit polyclonal anti-CRISPR-Cas9 antibody (ab191468, abcam, Cambridge, UK). Absence of specific bands at 163 kDa confirmed the absence of residual SpCas9 protein. Recombinant SpCas9 (rCas9) protein (62.5ng) (lane 1) was used as a positive control. **(B)** Whole cell lysate (5µg) from end of production UCB-TT52CAR19 GMP2 (UCB2) cells, untransduced (UnTD) or non-edited UCB-TT52CAR19 NO CAS9 GMP2 was analysed by EpiQuik CRISPR/Cas9 Assay ELISA Kit (Colorimetric) (EpigenTek, US) against manufacturer's standards (Std) and recombinant SpCas9 (rCas9) protein positive controls. SpCas9 protein levels in end of production UCB samples were comparable to blank control.

### Supplementary Figure S3.

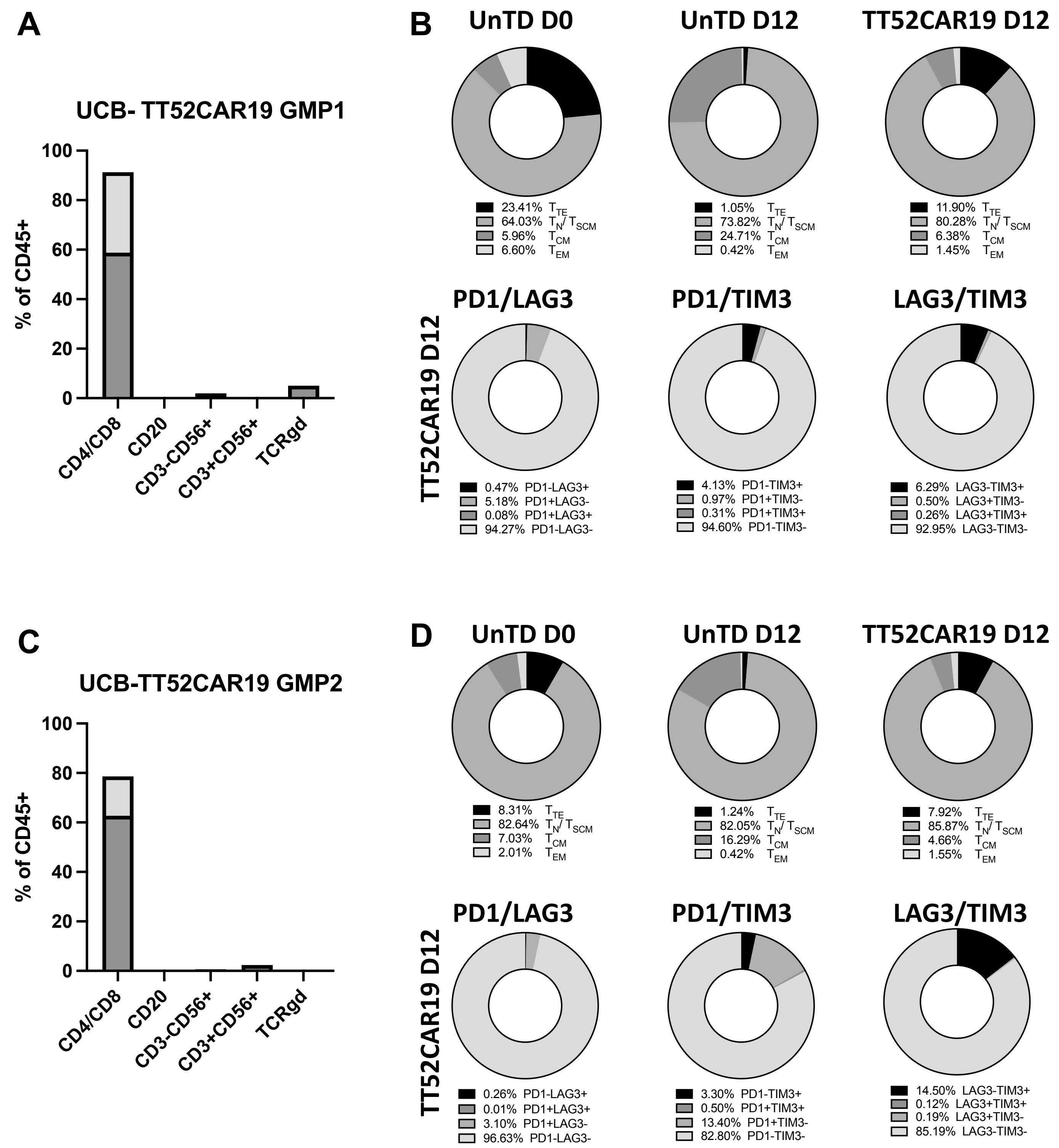


**Supplementary Figure S3. Memory phenotype of end of production UCB-TT52CAR19 GMP batches.** End of production UCB-TT52CAR19 cells, untransduced (UnTD) cells from day 0 (D0) and day 12 (D12) or UCB-TT52CAR19 cells from GMP batches GMP1 (UCB1) (top) or GMP2 (UCB2) (bottom) were phenotyped by flow cytometry. **(A, C)** Lymphocyte subset distribution of the two cell banks: T cells represented by combined CD4+ cells (dark grey) and $CD8^+$ (light grey); B cells expressing $CD20^+$; NK and T/NK cells ($CD3^-CD56^+$ and $CD3^+CD56^+$, respectively); TCRɣδ T cells; **(B top, D top)** T cell subsets exhibited surface markers $CD45RA^+CD62L^+$ attributed to naïve T cells (TN) and memory stem cell T cells (TSCM). Remaining subsets represented differentiated $CD45RA^-CD62L^+$ central memory (TCM), $CD45RA^+CD62L^-$ terminal effectors (TTE) and $CD45RA^-CD62L^-$ effector memory (TEM) T cells; **(B bottom, D bottom)** Profiling of activation/exhaustion of the three cell banks was investigated using three co-inhibitory markers (PD-1, TIM3, LAG3).

### Supplementary Figure S4.

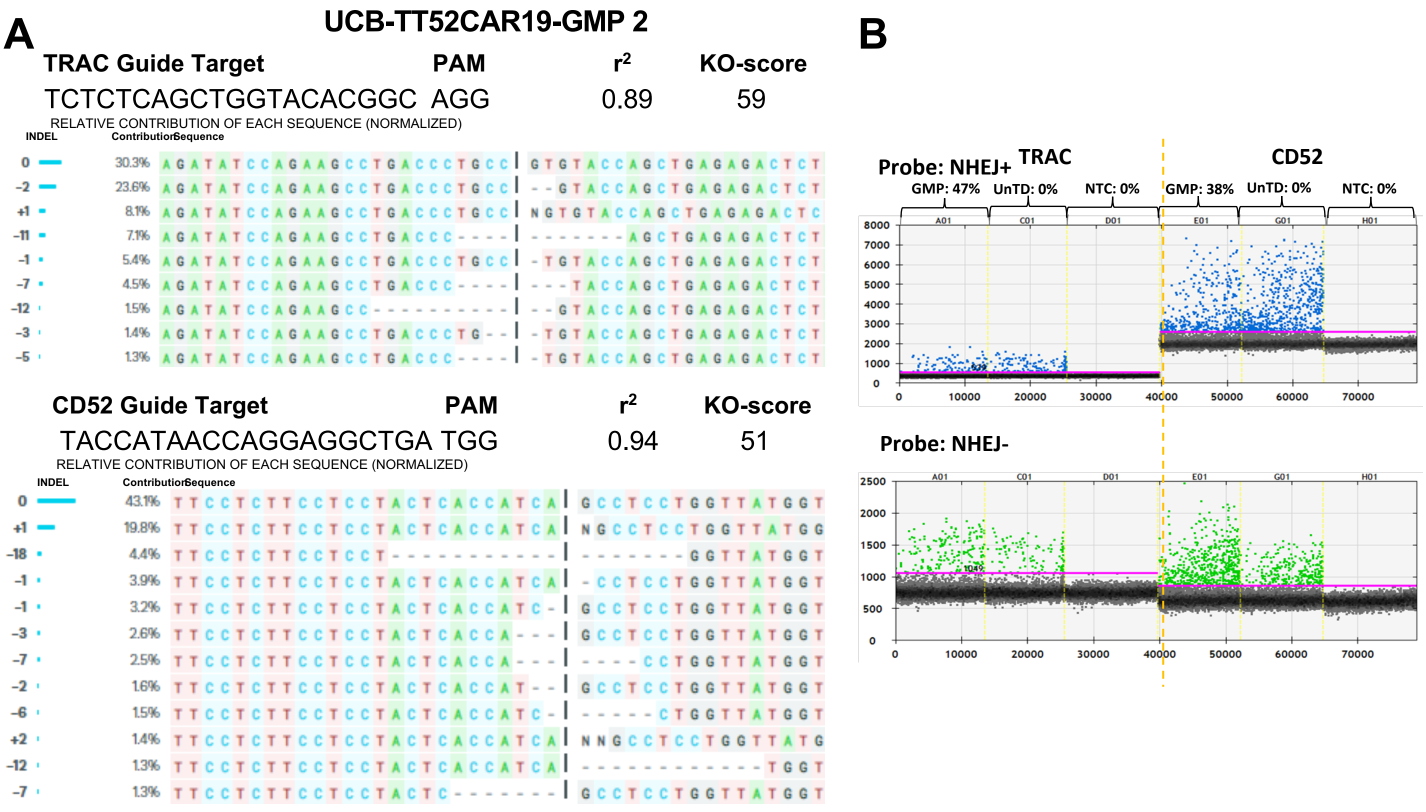


**Supplementary Figure S4. Molecular characterisation of on-target events following CRISPR-Cas9 mediated cleavage. (A)** ICE analysis of Sanger sequence traces identified indels in end of production UCB-TT52CAR19 GMP2 (UCB2) cells as signatures of NHEJ at the TRAC locus and at the CD52 target site. **(B)** Droplet Digital PCR (ddPCR) was used for the detection and quantification of indels by ddPCR at both the TRAC and CD52 sites using separate probes, one specific to the predicted NHEJ region (NHEJ$^{+}$) and a second outside the NHEJ region (NHEJ$^{-}$).

### Supplementary Figure S5.

#### Top 10 IS Gene Names, Frequencies and Location

| Rank | | Gene Name | Frequency [%] | Location |
|---|---|---|---|---|
| Top1 | | VIPR2 | 0.185 | 7+159068479 |
| Top2 | | NARF | 0.077 | 17+82456875 |
| Top3 | | TRAF2 | 0.068 | 9-136919938 |
| Top4 | | PTBP2 | 0.068 | 1+96752542 |
| Top5 | | PRRC2B | 0.06 | 9+131411295 |
| Top6 | | KIAA1715 | 0.057 | 2-175988556 |
| Top7 | | UQCC3 | 0.057 | 11-62673431 |
| Top8 | | MIR548AG1 | 0.057 | 4-59547877 |
| Top9 | | MAML2 | 0.057 | 11-96065506 |
| Top10 | | TNIK | 0.057 | 3-171093835 |
| #All Other mapp. IS | | 16853 | 99.257 | |

| | |
|---|---|
| Seq Count 10 Strongest | 880 |
| Seq Count all other mapp. IS | 117583 |
| Total Seq Counts Used | 118463 |

**Supplementary Figure S5. Integration site analysis.** Ligation-mediated PCR (LMPCR) detection and quantification of vector integration sites (IS) in UCB-TT52CAR19 GMP2 (UCB2) end of production cells where the top 10 most frequent sites comprised <0.1% of integrants.

### Supplementary Figure S6.

**A**

**UCB-TT52CAR19-GMP 2**

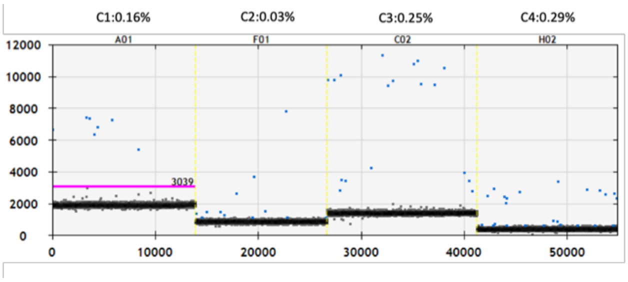


**B**

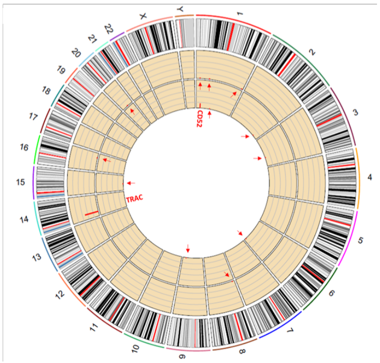


**TRAC**

| Index | Chromosome | ChromStart | ChromEnd | Mutant Frequency |
|---|---|---|---|---|
| TRAC-01 | chr14 | 23016465 | 23016487 | 50.58 |
| TRAC-02 | chr8 | 10627242 | 10627255 | 0.02 |
| TRAC-03 | chr1 | 68621798 | 68621815 | 0.11 |
| TRAC-04 | chr1 | 19612375 | 19612394 | 0.23 |
| TRAC-05 | chr1 | 242872248 | 242872256 | 0.02 |
| TRAC-06 | chr20 | 31622178 | 31622197 | 0.02 |
| TRAC-07 | chr16 | 66528021 | 66528030 | 0.05 |

**CD52**

| Index | Chromosome | ChromStart | ChromEnd | Mutant Frequency |
|---|---|---|---|---|
| CD52-01 | chr1 | 26644537 | 26644559 | 16.98 |
| CD52-02 | chr9 | 118431471 | 118431494 | 0.03 |
| CD52-03 | chr15 | 45497080 | 45497102 | 0.02 |
| CD52-04 | chr1 | 95875501 | 95875523 | 0.01 |
| CD52-05 | chr2 | 207739023 | 207739045 | 0.02 |
| CD52-06 | chr3 | 135905238 | 135905260 | 0.02 |
| CD52-07 | chr6 | 156954684 | 156954706 | 0.04 |

**Supplementary Figure S6. Molecular characterisation of off-target and predicted translocation events following CRISPR-Cas9 mediated cleavage. (A)** ddPCR based quantification of possible translocations after 'on-target' DNA scission. Low frequency translocation events (blue dots) C1-C4 arising between the edited TRAC and CD52 loci. Cumulative events for all four possible events were <1%. **(B)** Circos plot with verification of quantification using targeted NGS across six highest scoring predicted off-target sites. TRAC-01 (solid red line marking locus in outer yellow circle), CD52-01 (solid red line marking inner yellow circle) and at predicted off-target sites TRAC-02-TRAC-07 (red arrows marking outer yellow circle) and CD52-02-CD52-07 (red arrows marking inner yellow circle). Table shows that negligible off-target events were detected for both the TRAC and CD52 guides.

### Supplementary Figure S7.

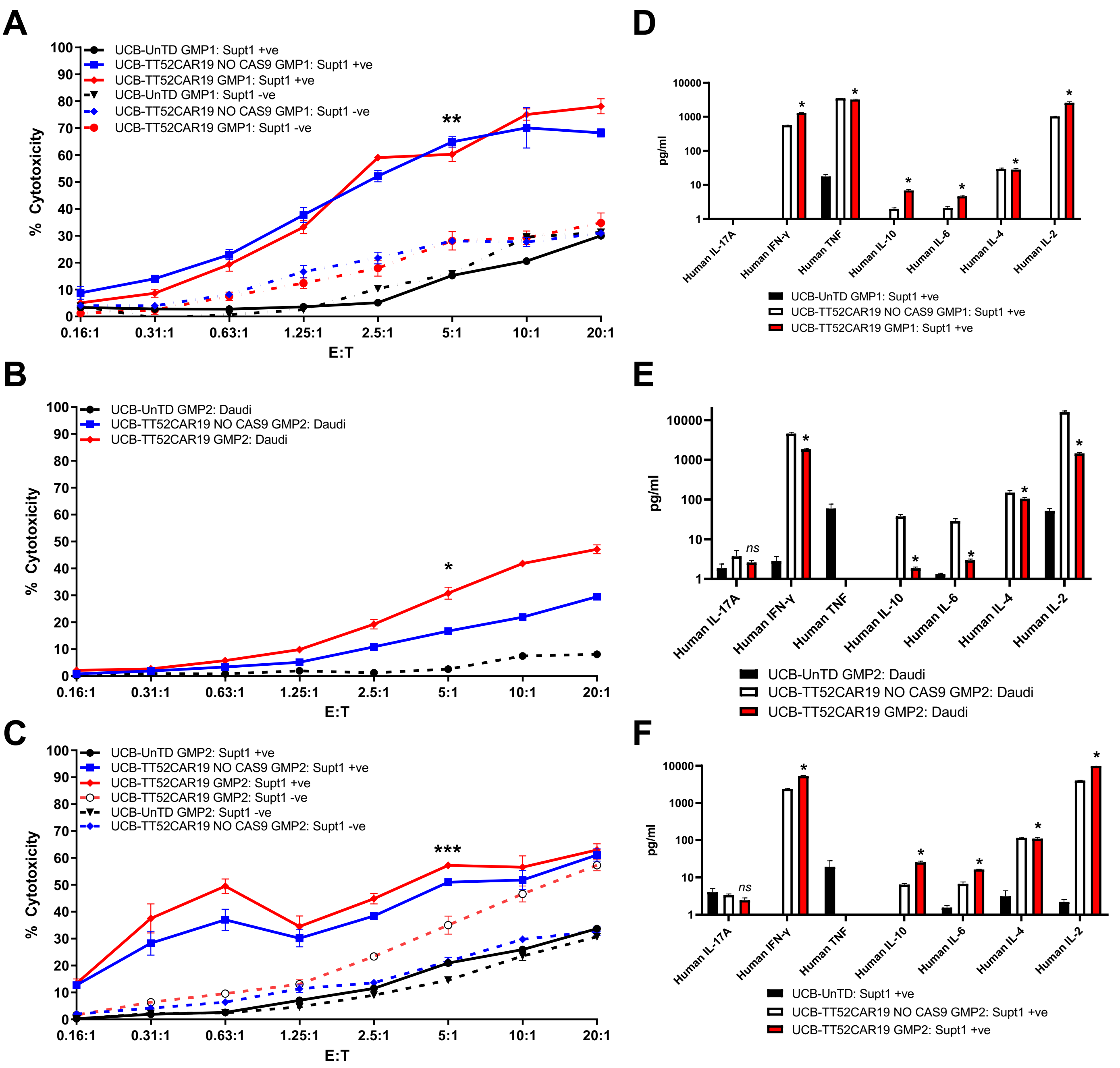


**Supplementary Figure S7. *In vitro* killing potential and cytokine production of UCB-TT52CAR19 cells against CD19⁺ targets.** In vitro cytotoxicity of UCB-TT52CAR19 cell banks compared to respective UCB-TT52CAR19 transduced but not edited (UCB-TT52CAR19 TCR$^+$) or non-transduced (UnTD) controls when measured by $^{51}$Cr chromium release of labelled Supt1 CD19$^+$ or CD19$^-$ cells **(A, C)** or CD19+ Daudi **(B)** cells following four hours co-culture at incremental effector (E) – target (T) ratio. Effector responses were considered successful if ≥50% lysis was detected compared to UnTD controls at E:T cell ratios between 1.25:1 and 5:1. Error bars SEM, n=3 replicate/wells. *P<0.05, **P<0.01, ***P<0.001 by t-test. In vitro target specific cytokine secretion of UCB-TT52CAR19 GMP cell banks, and respective UCB-TT52CAR19 TCR$^+$ or non-transduced (UnTD) controls after co-culture with Supt1 CD19$^+$ cells **(D, F)** or CD19$^+$ Daudi **(E)** cells overnight. The presence of cytokines in the co-culture supernatant was measured by cytokine bead array and levels >50pg/ml were considered positive responses. Significance was calculated between UCB-TT52CAR19 GMP1 and GMP2 banks and respective UnTD controls. *P<0.05 by t-test. Error bars represent SEM, n=3 replicates.

### Supplementary Figure S8.

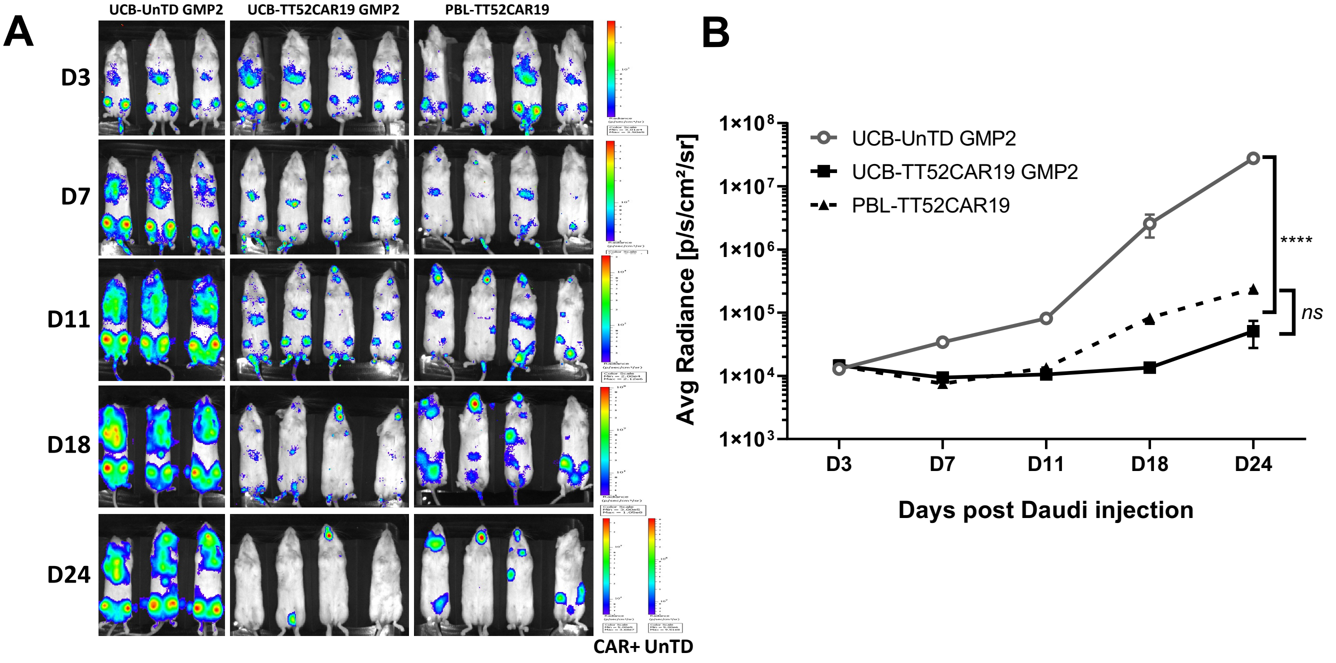


**Supplementary Figure S8. *In vivo* tumour clearance in human:murine xenograft model of CD19⁺ disease.**

Timeline of *in vivo* human:murine xenograft modelling indicating target and effector intravenous injection days and bioluminescent imaging (BLI) timepoints **(A)**. Serial measurement of bioluminescence of Daudi CD19⁺ B cell disease in immunodeficient mice NOD/SCID/γc (NSG) mice (n=12) infused with GFP/luciferase expressing Daudi CD19⁺ B cells were treated on day 4 with either umbilical cord blood (UCB) UCB-TT52CAR19 GMP2 (UCB2) (n=4) or peripheral blood lymphocyte (PBL) PBL-TT52CAR19 (n=4) and were monitored over a 4-week period **(B, C)** Non-transduced (UnTD) (n=3) T cells were used as controls. Error bars represent SEM. Significance compared by area under the curve using one-way ANOVA (F-value 10.63) with Tukey multiple comparison post-hoc; not significant (ns) $P \geq 0.05$; ****$P < 0.0001$.

### Table S1.

| Name | Application | Sequence | Primer/Probe |
|---|---|---|---|
| TRAC-on F | On- & off- targets NGS | CATGAGACCGTGACTTGCCA | Primer |
| TRAC-on R | On- & off- targets NGS | ACACATCAGAATCCTTACTTTGTGA | Primer |
| TRAC-02 F | On- & off- targets NGS | CAACTCTTGCTGCAACCTGA | Primer |
| TRAC-02 R | On- & off- targets NGS | TGCATCAGTCAACTTAGGTGAG | Primer |
| TRAC-03 F | On- & off- targets NGS | CCAAGATGGCAGAAGGGAAT | Primer |
| TRAC-03 R | On- & off- targets NGS | GAGGTCTTGCAAATTCAGGCT | Primer |
| TRAC-04 F | On- & off- targets NGS | AGCTTGAATGGCATCCTGAG | Primer |
| TRAC-04 R | On- & off- targets NGS | GCTTGGCCTCTCCAACTATG | Primer |
| TRAC-05 F | On- & off- targets NGS | CTGGAGGAAAGAAACAGACAGTAC | Primer |
| TRAC-05 R | On- & off- targets NGS | AAAAGGAGCCTGGCCATATTTC | Primer |
| TRAC-06 F | On- & off- targets NGS | CCGTCATCACACTGAACTTTGT | Primer |
| TRAC-06 R | On- & off- targets NGS | TCCCCATAACCTTTTCTGACCA | Primer |
| TRAC-07 F | On- & off- targets NGS | CACCCCAGCACCCACTATAC | Primer |
| TRAC-07 R | On- & off- targets NGS | GCCTCACAAGCAAGCCATAG | Primer |
| TRAC F | ddPCR NHEJ | TTGTCCATCACTGGCATC | Primer |
| TRAC R | ddPCR NHEJ | TGTGACACATTTGTTTGAGAATC | Primer |
| TRAC/NHEJ- | ddPCR NHEJ | TCATGTCCTAACCCTGATCCTCTTGT | Probe |
| TRAC/NHEJ+ | ddPCR NHEJ | ACCCTGCCGTGTACCAGCT | Probe |
| CD52-on F | On- & off- targets NGS | CAAGACAGCCACGAAGAT | Primer |
| CD52-on R | On- & off- targets NGS | AGGAGAGAAGGCTGGGTC | Primer |
| CD52-02 F | On- & off- targets NGS | AGCCAGCCTACTTGCCAA | Primer |
| CD52-02 R | On- & off- targets NGS | CCCAGCTTCTTAGGAGTGTC | Primer |
| CD52-03 F | On- & off- targets NGS | GTGTGTAGACTTCAAAGGGCA | Primer |
| CD52-03 R | On- & off- targets NGS | ACTGGGTAATTCTGAGTTGTGG | Primer |
| CD52-04 F | On- & off- targets NGS | CTGGTTTCCTTGGAGCCA | Primer |
| CD52-04 R | On- & off- targets NGS | GTAGACCTGAGCCACCTGAC | Primer |
| CD52-05 F | On- & off- targets NGS | TTGCCTTACCACTGAGCT | Primer |
| CD52-05 R | On- & off- targets NGS | AGACAAGTGCTGCCTTACCA | Primer |
| CD52-06 F | On- & off- targets NGS | CTAGATATCCATGGGTGATTGG | Primer |
| CD52-06 R | On- & off- targets NGS | GACCCAGTTCACTTCCTGCT | Primer |
| CD52-07 F | On- & off- targets NGS | ATGCAAGAGTGGCCCAAAT | Primer |
| CD52-07 R | On- & off- targets NGS | TCACTTGTTCTCCCCTGACC | Primer |
| CD52 F | ddPCR NHEJ | CAAGACAGCCACGAAGAT | Primer |
| CD52 R | ddPCR NHEJ | AGGAGAGAAGGCTGGGTC | Primer |
| CD52/NHEJ- | ddPCR NHEJ | CCAAAGTTGCTTGGCATGGA | Probe |
| CD52/NHEJ+ | ddPCR NHEJ | CATCAGCCTCCTGGTTATGGTACA | Probe |
| TRAC F + CD52 R | ddPCR Translocations | TTGTCCATCACTGGCATC | Primer F |
| TRAC F + CD52 R | ddPCR Translocations | GCCAGGCGTTGCTCTTAC | Primer R |
| CD52 F + TRAC R | ddPCR Translocations | CAAGACAGCCACGAAGAT | Primer F |
| CD52 F + TRAC R | ddPCR Translocations | TGTGACACATTTGTTTGAGAATC | Primer R |
| TRAC R + CD52 R | ddPCR Translocations | TCAGAATCCTTACTTTGTGACAC | Primer F |
| TRAC R + CD52 R | ddPCR Translocations | GCCAGGCGTTGCTCTTAC | Primer R |
| CD52 F + TRAC F | ddPCR Translocations | CAAGACAGCCACGAAGAT | Primer F |
| CD52 F + TRAC F | ddPCR Translocations | ATCACTGGCATCTGGACTC | Primer R |
| TRAC F + CD52 R | ddPCR Translocations | TCATGTCCTAACCCTGATCCTCTTGT | Probe |
| CD52 F + TRAC R | ddPCR Translocations | AGTCTGTCTGCCTATTCACCGATT | Probe |
| TRAC R + CD52 R | ddPCR Translocations | TCGGTGAATAGGCAGACAGACTTGT | Probe |
| CD52 F + TRAC F | ddPCR Translocations | TGGGACAAGAGGATCAGGGT | Probe |
| Albumin F | ddPCR Translocations | GCTGCTATCTCTTGTGGGCTGT | Primer |
| Albumin R | ddPCR Translocations | ACTCATGGGAGCTGCTGGTTC | Primer |
| Albumin | ddPCR Translocations | CCTGTCATGCCCACACAAATCTCTCC | Probe |

**Table S1. On-, off-target and translocation primer and probe sets.**
Primer and probe sets used for quantification of ‘on-target’ and ‘off-target’ NHEJ events or possible translocations after ‘on-target’ DNA scission in UCB-TT52CAR19 Good Manufacturing Process (GMP) batches GMP1 (UCB1) and GMP2 (UCB2).
